# CONFOLD2: Improved contact-driven ab initio protein structure modeling

**DOI:** 10.1101/228460

**Authors:** Badri Adhikari, Jianlin Cheng

## Abstract

**Background:** Contact-guided protein structure prediction methods are becoming more and more successful because of the latest advances in residue-residue contact prediction. To support the contact-driven structure prediction, effective tools that can quickly build tertiary structural models of good quality from predicted contacts need to be developed.

**Results:** We develop an improved contact-driven protein modeling method, CONFOLD2, and study how it may be effectively used for ab initio protein structure prediction with predicted contacts as input. It builds models using various subsets of input contacts to explore the fold space under the guidance of a soft square energy function, and then clusters the models to obtain top five models. CONFOLD2 is benchmarked on various datasets including CASP11 and 12 datasets with publicly available predicted contacts and yields better performance than the popular CONFOLD method.

**Conclusion:** CONFOLD2 allows to quickly generate top five structural models for a protein sequence, when its secondary structures and contacts predictions at hand. CONFOLD2 is publicly available at https://github.com/multicom-toolbox/CONFOLD2/.

## Background

The most successful ab initio protein structure methods, i.e. fragment-assembly based methods, require generating a lot of decoys to deliver accurate predictions. Methods that can build models faster and are more residue contact sensitive are needed to realize the promise of ab initio protein structure prediction driven by the recent advances in contact prediction [1,2]. The CONFOLD method [3] can build high quality secondary structures (including beta-sheets) and correct tertiary structures when predicted contacts are accurate. It is integrated into other protein structure prediction methods like CoinFold [4] and PconsFold2 [2]. In this paper, we develop an improved version of CONFOLD by incorporating a soft-square energy function into CONFOLD, building models using multiple sub-sets of contacts, adding model selection capability, and rigorously testing it on various datasets including the Critical Assessment of protein Structure Prediction (CASP) 11 and 12 datasets. CONFOLD2 also addresses a major limitation of the CONFOLD method, i.e. generating a decoy of 200 models and not producing top one or top five models. Compared to fragment-assembly methods that need to generate thousands of model decoys [5], CONFOLD2 explores the fold space by generating just a few hundred model decoys, and hence it runs relatively fast.

## Implementation

Recently, it is found that energy functions that do not penalize unsatisfied predicted contacts after certain distance threshold yield more accurate model reconstruction [5-7]. Different contact energy functions like FADE [5], square-well function with exponential decay [6], and modified Lorentz potential [7] applied to contact-guided protein folding have been found to work best for various folding algorithms, mostly fragment-assembly based methods. When distance geometry based approaches are used to fold proteins with restraints, it has been shown that soft-square function performs best, with the ‘rswitch’ parameter to be tuned [8]

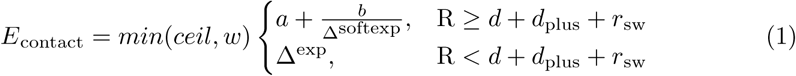

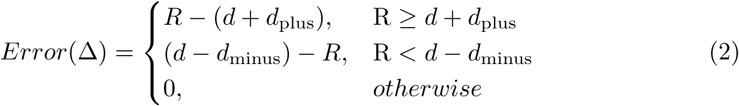

We replaced CONFOLD’s [3] soft-square asymptotic energy function (designed originally for the experimental NOE restraints) with the soft-square function (**Equation (1)**), where the error is defined in **Equation (2)**. The parameter d, d_minus_,and d_plus_ define the interval [d-d_minus_, d+d_plus_], where the error is zero. For contacts predicted to be less than 8Å distance, we set d, d_minus_, and d_plus_ to 3.6, 0.1, and 4.4 respectively. The switching parameter r_sw_ defines the boundary where the square error function starts to taper into a constant error (see **Figure1**). R is the actual distance between C*β* atoms of the predicted contact residue pair in the model. The exponents, ‘exp’ and ‘softexp’ are both set to 2. Since the contact weight multiplies the energy term, the maximum weight (ceil) that any pair of predicted contacts can have is set to 1000, and ‘w’ is the weight of each contact pair and is set to 1. The most important parameter affecting the quality of reconstruction is r_sw_ and we optimized it to be 1.8. ‘a’ and ‘b’ are constants determined at run-time such that the function is smooth at r_sw_ equal to 1.8. Our soft-square contact energy term is calculated either using a square error function or approximately constant error function based on a switching parameter - r_sw_. It defines a threshold until which the error increases as a square error function and beyond which the error tapers to a constant error. **Figure1** demonstrates how the switching parameter affects the overall energy calculations.

Using the soft-square function as contact energy term, CONFOLD2 initially predicts 200 models using various subsets of input contacts, and selects five top models by clustering them. To effectively explore the fold space captured by the predicted contacts, we prepare 40 different subsets of input contacts by selecting top xL contacts, where x = 0.1, 0.2, 0.3, …, 4.0 and L is length of the protein, and build 20 models for each subset. For each of this subset of contacts, top 5 models in the second stage of CONFOLD modeling are selected based on the contact energy score, resulting in a total of 200 models. Next, to filter out unfolded models, we rank these 200 models by calculating their contact satisfaction score using top L/5 long-range contacts, and filter out the bottom 150 models. The remaining 50 models are clustered into five clusters by calculating their pairwise structural similarity measured by TM-score. We select the five models closest to the centroids of these five clusters as the top five predictions with the rank determined by the satisfaction score of the top L/5 long-range contacts. SCRATCH suite [9] is used to predict three-state secondary structure and Maxcluster [10] to compute pairwise model similarity for clustering.

**Figure 1.**
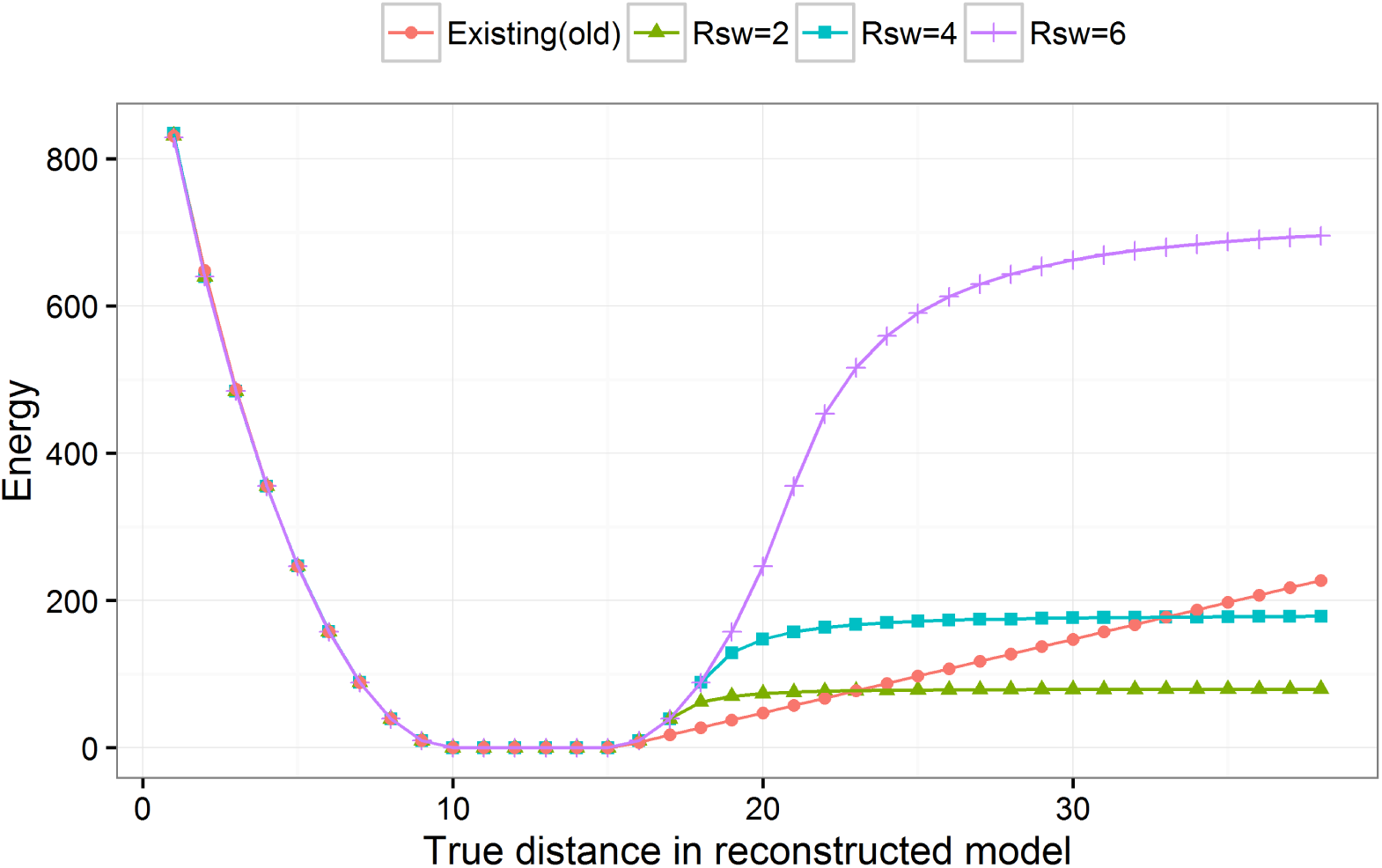
Behavior of the contact energy term for various r_sw_ values. For this demonstration desired distance is set to 10 Å with a lower-bound of 0 Åand upper-bound of 5 Å, i.e. the desired distance between the pair of restrained residues is 10.0 Å and 15.0 Å. The “Existing" energy calculations refers to the old energy term implemented in CONFOLD method. The plot shows that depending upon the switching parameter, r_sw_, the energy calculations can taper early at around 1 or 2 Å for r_sw_ = 2 or at more than 25 Å for r_sw_ = 6.

## Results

As the first benchmark, we compared the performance of CONFOLD2 with the original CONFOLD method [3] on the 150 proteins in the PSICOV dataset [11] using the contacts predicted using PSICOV [11] (see **Table1**). The original CONFOLD method generates top 200 models and provides no ranking of the reconstructed models, so we compare the two methods using best-of-200 models. On the PSICOV dataset, when best of 200 models are evaluated, CONFOLD2 achieves a mean TM-score of 0.57 compared to 0.55 of CONFOLD. This improvement in CONFOLD2 is statistically significant per paired t-test with a p-value of 4 x 10^-8^ (see **Suppl. Table S1** for a detailed comparison).

Next, to evaluate our model selection technique (selecting top five models from 200) we compared our approach of model selection using clustering with the model ranking using contact satisfaction score only. On the same dataset, when we selected top five models using contact satisfaction score of top L/5 or L/2 long-range contacts, we achieved best-of-top-five TM-score of 0.50. The rationale for using top L/5 or L/2 contacts (instead of L or more) is that these subsets are found to best reflect the accuracy of the predicted contacts [12]. In contrast, when we filter out the bottom 150 models, cluster the remaining 50 into five clusters, and select the cluster centroids, we obtain best-of-top-five TM-score of 0.52, suggesting that the clustering approach is effective in selecting models built from contacts. As summarized in **Table1**, we also reconstructed models for the PSICOV-150 dataset using contacts predicted by MetaPSICOV [13] and obtained a mean TM-score of 0.62 when best of top-five models are evaluated (see **Supp. Table S1** for detailed results), indicating that the improved contact prediction leads to the better tertiary structure reconstruction.

Finally, using CONFOLD2, we predicted models for the protein sequence targets in CASP11 and CASP12 datasets with contacts predicted by the most accurate predictor in each of the CASP experiments – CONSIP2 [14] in CASP11 and Raptor-X [15] in CASP12 (see **Table1**). The average TM-score of the reconstructed models for both CASP11 and 12 datasets is 0.46 when best-of-200 models are evaluated and 0.41 when best-of-five models are evaluated.

**Table 1.** Summary of the performance of CONFOLD2 on PSICOV, CASP11, and CASP12 datasets. Mean contact precision of top L/5 for (i) all (short-range, medium-range, and long-range: P_SR+MR+LR_) contacts, and (ii) long-range contacts (P_LR_) is reported for all the datasets. The TM-score of the best-of-200 and best-of-5 models reconstructed by CONFOLD2 are also presented. Results for single-domain and multi-domain subsets of the CASP11 and CASP12 datasets are also reported separately.

| Dataset | Contact Precision (L/5) |  |  | TM-score of Models |  |
| --- | --- | --- | --- | --- | --- |
| | Method | $P_{SR+MR+LR}$ | $P_{LR}$ | Best-of-200 | Best-of-5 |
| PSICOV-150 | PSICOV | 72.6 | 64.0 | 0.57 | 0.52 |
| PSICOV-150 | MetaPSICOV | 88.4 | 77.2 | 0.65 | 0.62 |
| CASP12 All | Raptor-X | 71.3 | 58.6 | 0.46 | 0.41 |
| CASP12 Single Domain | Raptor-X | 70.6 | 58.6 | 0.49 | 0.44 |
| CASP12 Multi-Domain | Raptor-X | 72.0 | 58.7 | 0.44 | 0.38 |
| CASP11 All | CONSIP2 | 71.8 | 50.2 | 0.46 | 0.41 |
| CASP11 Single Domain | CONSIP2 | 75.8 | 57.4 | 0.52 | 0.48 |
| CASP11 Multi-Domain | CONSIP2 | 67.7 | 42.4 | 0.40 | 0.34 |

## Discussion

Observing the lower reconstruction accuracy for the CASP datasets compared to the PSICOV dataset, we investigated if the performance was affected by multi-domain proteins because we build models for the whole targets first and evaluated them at domain level. As shown in **Table1**, the reconstruction accuracy is higher for single domain proteins than multi-domain proteins (see **Supp. Table S2 and S3** for details). Yet, the reconstruction accuracy for single domain proteins is still lower than that of the PSICOV dataset. For the further investigation, from the single domain proteins in both CASP11 and 12 datasets, we removed some proteins with low accuracy contact predictions so that both datasets have the mean contact precision of top L/5 long-range contacts the same as that of the PSICOV dataset, i.e. precision = 64%. On such reduced datasets, the average TM-scores of the best-of-200 models for CASP11 and 12 proteins are 0.55 and 0.52 respectively, which are slightly lower than the mean TM-score for PSICOV dataset (0.57). Since TM-score of 0.5 is the threshold if the topology of a protein structure is correctly predicted, for all three datasets, it can be concluded that the fold of single domain proteins can be reconstructed correctly (TM-score ≥ 0.5) on average if the precision of predicted long-range contacts is at least 64%. Although the sequence lengths of the domains in the CASP datasets are much higher than the PSICOV-150 dataset, which have up to 500 residues, we did not find any substantial correlation between the domain length and the reconstruction accuracy.

## Conclusions

We have developed CONFOLD2, a method for building three-dimensional protein models using predicted contacts and secondary structures. It explores the fold space captured in predicted contacts by creating various subsets of predicted contacts and builds decoy sets, and then clusters the decoys to obtain top five models. CONFOLD2 is significantly better than the original CONFOLD method. Structure predictions using some recently available contact prediction datasets, show that the for most protein sequences CONFOLD2 is able to capture the structural fold of the protein.

## Availability and requirements

Project name: CONFOLD2

Project home page: https://github.com/multicom-toolbox/CONFOLD2

Operating systems: Platform independent

Programming language: Perl

Other requirements: Perl interpreter, CNS suite, TM-score (included), MaxCluster (included), and DSSP (included)

License: GNU GPL

Any restrictions to use by non-academics: None

## Acknowledgements

Some of the computation for this work was performed on the high performance computing infrastructure provided by Research Computing Support Services and in part by the National Science Foundation under grant number CNS-1429294 at the University of Missouri, Columbia MO). We would like to thank the RCSS team for their infrastructure and technical support.

## Additional Files

**Additional file 1 — Supplementary Tables**

Docx file containing various tables with detailed results.

